# Three-dimensional convolutional autoencoder extracts features of structural brain images with a “diagnostic label-free” approach: Application to schizophrenia datasets

**DOI:** 10.1101/2020.08.24.213447

**Authors:** Hiroyuki Yamaguchi, Yuki Hashimoto, Genichi Sugihara, Jun Miyata, Toshiya Murai, Hidehiko Takahashi, Manabu Honda, Akitoyo Hishimoto, Yuichi Yamashita

## Abstract

There has been increasing interest in performing psychiatric brain imaging studies using deep learning. However, most studies in this field disregard three-dimensional (3D) spatial information and targeted disease discrimination, without considering the genetic and clinical heterogeneity of psychiatric disorders. The purpose of this study was to investigate the efficacy of a 3D convolutional autoencoder (CAE) for extracting features related to psychiatric disorders without diagnostic labels. The network was trained using a Kyoto University dataset including 82 patients with schizophrenia (SZ) and 90 healthy subjects (HS), and was evaluated using Center for Biomedical Research Excellence (COBRE) datasets including 71 SZ patients and 71 HS. The proposed 3D-CAEs were successfully reconstructed into high-resolution 3D structural magnetic resonance imaging (MRI) scans with sufficiently low errors. In addition, the features extracted using 3D-CAE retained the relevant clinical information. We explored the appropriate hyper parameter range of 3D-CAE, and it was suggested that a model with eight convolution layers might be relevant to extract features for predicting the dose of medication and symptom severity in schizophrenia.

## Introduction

Deep learning (DL) has dramatically improved technology in speech recognition, image recognition, and many other fields^1^. Medical imaging can benefit greatly from recent progress in image classification and object detection using this cutting-edge technology^1^. In particular, as the global burden of psychiatric disorders increases^3,4^, psychiatric brain imaging studies using DL are anticipated to bring much benefit to society^5^. There are two major concerns about applying DL to psychiatric brain imaging: (1) treatment of the high dimensionality of data, and (2) the heterogeneity of psychiatric disorders^6^.

The dimensionality of raw magnetic resonance imaging (MRI) data is very high (often running into the millions) and large computer resources are required to analyze them. In order to reduce computational demands, in most neuroimaging studies several feature extraction methods have been used. Region of interests (ROIs), one of the most popular methods of feature extraction, has contributed to the detection of various structural and functional abnormalities in the brains of patients with psychiatric disorders^7-10^. ROIs (often dozens or hundreds) are usually set based on neuroscience knowledge^11^. For example, average gray matter volumes or cortical thicknesses at specific ROIs are extracted as features and then the relationship between the features and disease clinical information is analyzed^12-14^. Even in the studies using DL, ROI-based features are often used as input^3,15,16^. In addition, many DL studies avoid using high-resolution three-dimensional (3D) images directly, but instead DL networks are trained using two-dimensional slices^3,17,18^. A limitation of these studies is that they ignore the 3D spatial information contained within the original MRI scans.

In recent years, with improvements in computer performance and refinement of computational techniques, studies have investigated the ways to treat high-resolution 3D MRI scans as inputs to DL. For example, Wang, et al.^19^ successfully discriminated Alzheimer’s dementia from healthy subjects using high-resolution 3D MRI data as input to DL. Similar attempts have been made for discriminating psychiatric disorders including schizophrenia^20^ and developmental disorders^21^. Although these studies demonstrated that DL can be applicable to the analysis of high-resolution 3D MRI data, discrimination-based approaches may be challenging due to the heterogeneity of psychiatric disorders.

Heterogeneity is one of the main challenges that current psychiatric research faces^6^. The current symptom-based definitions of psychiatric disorders, standardized in the Diagnostic and Statistical Manual of the American Psychiatric Association (DSM)^22^ and the International Classification of Diseases (ICD)^23^, have been highlighted as lacking predictive and clinical validity due to genetic and clinical heterogeneity^24^. For example, in schizophrenia, a recent study found evidence for significant overlapping of the relatively common risk variants that are tagged in genome-wide association studies (GWAS) of between several psychiatric disorders, and there may also be lower genetic correlation within disorders^25^. In addition, even in patients given the same diagnosis of schizophrenia, the severity of symptoms, response to medication, and prognosis often vary widely among patients^26,27^. Therefore, in psychiatric disorders research, a simple competition for discrimination accuracy based on the current disorder categories may be insufficient to elucidate on pathophysiology, although most current studies using DL are attempting to discriminate disease in healthy subjects^3,28^.

One possible alternative direction for using DL techniques in psychiatric neuroimaging studies may be for diagnostic label-free feature extraction. In the current study, we focus on an autoencoder (AE) as a DL algorithm that allows feature extraction without labels^29^. Indeed, there are some studies that have used AE-based feature extraction for psychiatric neuroimaging. For example, Pinaya et al. extracted features from structural MRI scans using AE, i.e., without using diagnostic labels. The authors successfully predicted the age and gender of participants, and discriminated patients with autism spectrum disorders (ASD) and schizophrenia from healthy subjects^16^. However, these studies used ROI-based features such as cortical thickness and functional connectivity as inputs to the AE. As such, the use of high-resolution 3D brain images for inputs to the AE remains challenging, with a few exceptions. For example, Martinez-Murcia et al. extracted features from high-resolution 3D brain MRI data of patients with Alzheimer’s dementia using a 3D convolutional autoencoder (3D-CAE)^30^, they demonstrated that extracted features were useful for predicting age and Mini-Mental State Examination (MMSE) scores. This supports the efficacy of labeling free features based on 3D-CAE with high-resolution MRI. However, particularly when investigating psychiatric disorders, the appropriate architecture of 3D-CAE has not been fully investigated.

The purpose of this study was to investigate an efficient 3D-CAE-based feature extraction for the neuroimaging of psychiatric disorders. More specifically, in the current study, we used datasets that included patients with schizophrenia, a condition that has frequently been reported to be heterogeneous in previous neuroimaging studies^31^. The key points of our study are: (1) to use high-resolution 3D MRI data while preserving spatial information, and (2) diagnostic label-free feature extraction using 3D-CAE. For this purpose, we explored appropriate network structures of 3D-CAE by comparing the relationships between the features extracted by the model with different network structures and varying clinical information.

## Methods

### Experimental overview

Figure 1 illustrates an experimental overview of our study. We used two datasets, including participants diagnosed with schizophrenia as well as healthy subjects: a dataset collected at Kyoto University (Kyoto dataset) and a public dataset, The Center for Biomedical Research Excellence (COBRE; http://fcon_1000.projects.nitrc.org/indi/retro/cobre.html) dataset. (1) Gray matter was first extracted from the structural MRI data as preprocessing. (2) We then trained 3D-CAE to extract a latent feature representation from structural MRI using the Kyoto dataset. Sixteen 3D-CAEs with varying network structures were prepared for investigation of the optimal network depth and complexity. (3) Subsequently, the COBRE dataset was used to evaluate the applicability to another dataset. (4) Finally, we evaluated whether the extracted features retained clinical information by linear regression of the clinical information using the COBRE dataset.

**Figure 1.**
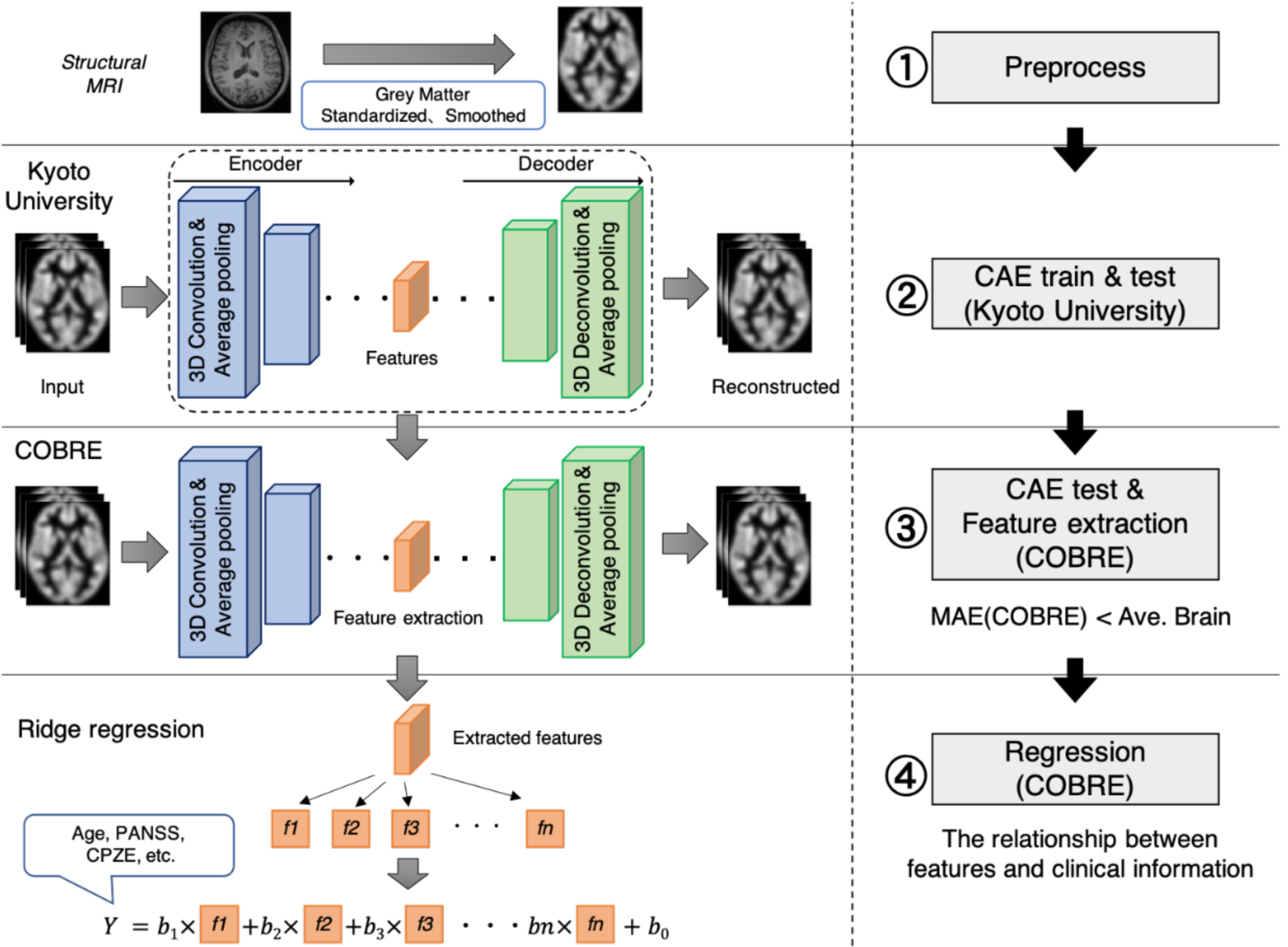
Experimental overview. 1. Preprocessing: The gray matter was extracted from the structural MRI, and standardized and smoothed using SPM. 2. CAE training: A schematic diagram is shown. 3D images of the Kyoto dataset were input, features were extracted, and the original image was reconstructed. 3. Feature extraction: the model trained using the Kyoto dataset was adopted to the COBRE dataset without updating the weights. 4. Linear regression: Each extracted feature was an explanatory variable, and demographic and clinical information were objective variables. Regression errors were evaluated, and the relationship between features and demographic and clinical information was investigated. 3D, three-dimensional; CAE, convolutional autoencoder; COBRE, Center for Biomedical Research Excellence; CPZE, dose of antipsychotic medication; MRI, magnetic resonance imaging; SPM, Statistical Parametric Mapping.

### Kyoto dataset description

A total of 172 subjects were investigated in this study, including 82 patients with schizophrenia and 90 healthy subjects. Patients were recruited from hospitals in Kyoto, Japan, and diagnosed by psychiatrists using the Diagnostic and Statistical Manual of Mental Disorders, 4^th^ edition (DSM-IV) ^32^ criteria for schizophrenia, confirmed with the patient edition of the Structured Clinical Interview for DSM-IV Axis I Disorders (SCID)^33^. No patients had any comorbid DSM-IV Axis I disorder. The clinical symptoms of all patients were estimated using the Positive and Negative Syndrome Scale (PANSS)^34^. Healthy subjects were screened with the non-patient edition of the SCID, confirming no history of psychiatric disorders. Exclusion criteria for all individuals included a history of head trauma, neurological illness, serious medical or surgical illness, or substance abuse. Note that participants were already diagnosed in order to expedite the data collection, but the diagnostic labels were not used to train the networks. All study participants signed an informed consent form. The study was performed in accordance with the current Ethical Guidelines for Medical and Health Research Involving Human Subjects in Japan and was approved by the Committee on Medical Ethics of Kyoto University and National Center of Neurology and Psychiatry.

All participants were scanned with a 3.0-Tesla Siemens Trio scanner (Siemens Healthineers, Erlangen, Germany). The scanning parameters of the T1-weighted 3D magnetization-prepared rapid gradient-echo (3D-MPRAGE) sequences were as follows: echo time (TE) = 4.38 ms; repetition time (TR) = 2,000 ms; inversion time (TI) = 990 ms; field of view (FOV) = 225 mm × 240 mm; acquisition matrix size = 240 × 256 × 208; resolution = 0.9375 × 0.9375 × 1.0 mm^3^.

### COBRE dataset description

In this study, the COBRE dataset, which is a public dataset, was acquired as a dataset with different scanning sites and parameters to the Kyoto University dataset. All the subjects were diagnosed on and screened with the SCID. The clinical symptoms of all patients were estimated using the PANSS. Exclusion criteria for individuals included a history of head trauma, neurological illness, serious medical or surgical illness, or substance abuse. We included a total of 142 subjects from this database in our study, including 71 patients with schizophrenia and 71 healthy subjects. As stated earlier, the diagnostic labels were not used to train the network.

MRI data were acquired using a 3.0-Tesla Siemens Tim Trio scanner (Siemens Healthineers). The scanning parameters of the T1-weighted 3D-MPRAGE sequences were as follows: TE = 1.64 ms; TR = 2,530 ms; TI = 900 ms; FOV = 256 mm × 256 mm; acquisition matrix size = 256 × 256 × 176; resolution = 1.0 × 1.0 × 1.0 mm^3^.

### Division of train, validation, and test

The 3D-CAE was trained using the Kyoto dataset. The dataset was randomly partitioned into training data, validation data, and test data (138 subjects, 16 subjects and 18 subjects, respectively). Training data, validation data, and test data were used for the training of the 3D-CAE, the validation of the model during training, and the final evaluation of generalizability within the datasets independent of the training and validation data, respectively. The COBRE dataset (142 subjects) was also used to evaluate the applicability of the network to another dataset.

The regression was carried out using the COBRE dataset. The five-fold cross validation technique was applied. Namely, the COBRE dataset samples (142 subjects) were randomly divided into five subgroups (four groups for training and one group for validation) and cross-validated by changing the combinations of groups. This five-fold cross-validation process was repeated ten times. Note that only patients with schizophrenia had clinical information available for analysis, and regressions based on the clinical information were performed using data from patients with schizophrenia (71 subjects). The details for the division of data are shown in Table 1.

**Table 1.**
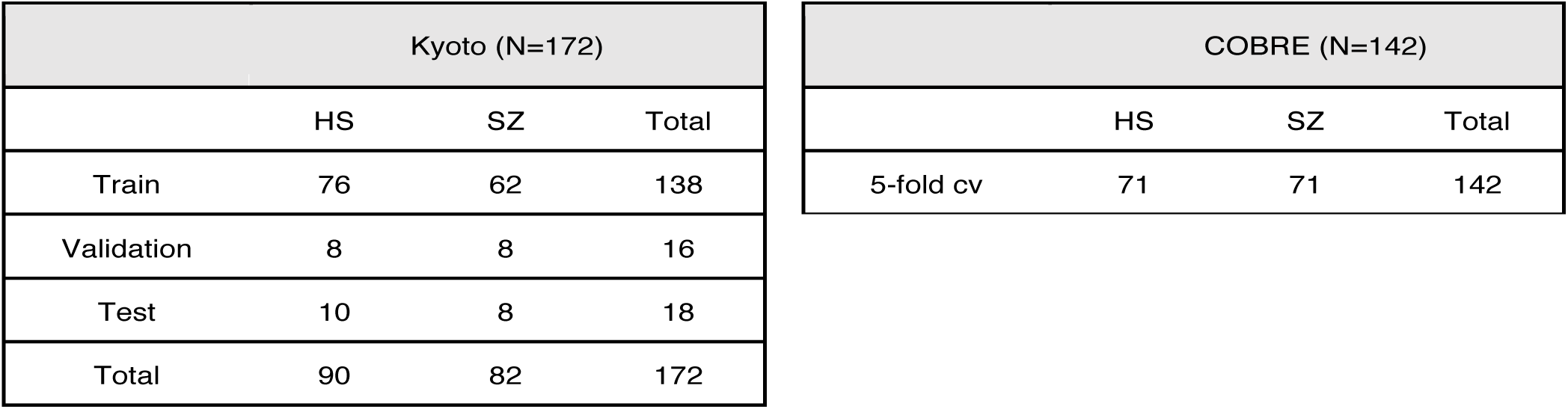
Division of dataset. The Kyoto dataset was used to develop the 3D-CAE model and was divided into train, validation and test dataset. The COBRE dataset was prepared for verification. At the time of regression, 5-fold-cross validation was performed.

| Kyoto (N=172) |  |  |  |
| --- | --- | --- | --- |
|  | HS | SZ | Total |
| Train | 76 | 62 | 138 |
| Validation | 8 | 8 | 16 |
| Test | 10 | 8 | 18 |
| Total | 90 | 82 | 172 |

### MRI preprocessing

The preprocessing was conducted using Statistical Parametric Mapping (SPM12, Wellcome Department of Cognitive Neurology, London, UK; https://www.fil.ion.ucl.ac.uk/spm/software/spm12/)^35^ with the Diffeomorphic Anatomical Registration Exponentiated Lie Algebra (DARTEL) registration algorithm^36^. All of the T1 whole brain structural MRI scans were segmented into gray matter (GM), white matter, and cerebrospinal fluid. Individual GM images were normalized to the standard Montreal Neurological Institute (MNI) template with a 1.5 × 1.5 × 1.5 mm^3^ voxel size and modulated for GM volumes. All normalized GM images were smoothed with a Gaussian kernel of 8 mm full width at half maximum (FWHM). Subsequently, each image was cropped to remove background as much as possible. The GM area was extracted from original images using a binary mask, created using SPM12. As a result, the size of input images to the 3D-CAE was 121 × 145 × 121 voxels.

Subsequently, the range of signal intensities in each image was normalized with a mean of 0 and a standard deviation of 1. The standardized value of voxel *i* in the sample *s, x*′_*s,i*_, was calculated as follows:

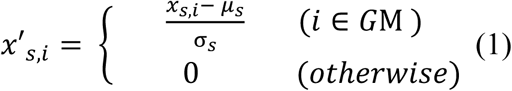

where *x*_*s,i*_ is the original value of intensity. *μ*_*s*_ and σ_*s*_,were average and standard deviation of all voxels contained in the GM area of sample *s*, respectively.

### Convolutional autoencoder training

An autoencoder is a kind of DL consisting of the encoder and the decoder. The encoder learns latent representations and reduces the dimension of the input. The decoder learns to reconstruct the input as close as possible to the original using the latent representations. 3D-CAE extends this architecture by using convolutional layers that can extract features directly from 3D images^37-39^. The CAE has two main hyper parameters: the number of convolutional layers and the number of channels, which are the target of the current study.

The convolutional layers apply a filter to an input to create feature maps that summarizes the features detected in the input. The feature maps are created for the number of channels. Since the convolutional layer generates feature maps while capturing the spatial information of the matrix, convolutional neural networks are beneficial to learning features of images. As the number of channels increases, the complexity of a model increases. Also, as the number of convolutions increases, the effective receptive field increases, thus allowing global and abstract features to be extracted. The effective receptive field is a region of the original image that can potentially influence the activation of neurons^40,41^.

The impact of two hyper parameters, the number of convolutional layers, and the number of channels was investigated. As shown in Figure 2, the set of two convolution/deconvolution layers and one pooling/unpooling layer was defined as a convolution/deconvolution “block”. In this experiment, the number of blocks was set ranging from 1 block to 4 blocks. The number of channels in the extraction layer was varied with 1, 4, 16, and 32 channels, but, the number of channels for other layers were fixed at 32. As a result, we created sixteen 3D-CAE models (4 block conditions × 4 channel conditions) to explore the effective range of hyper parameters for psychiatric brain imaging.

**Figure 2.**
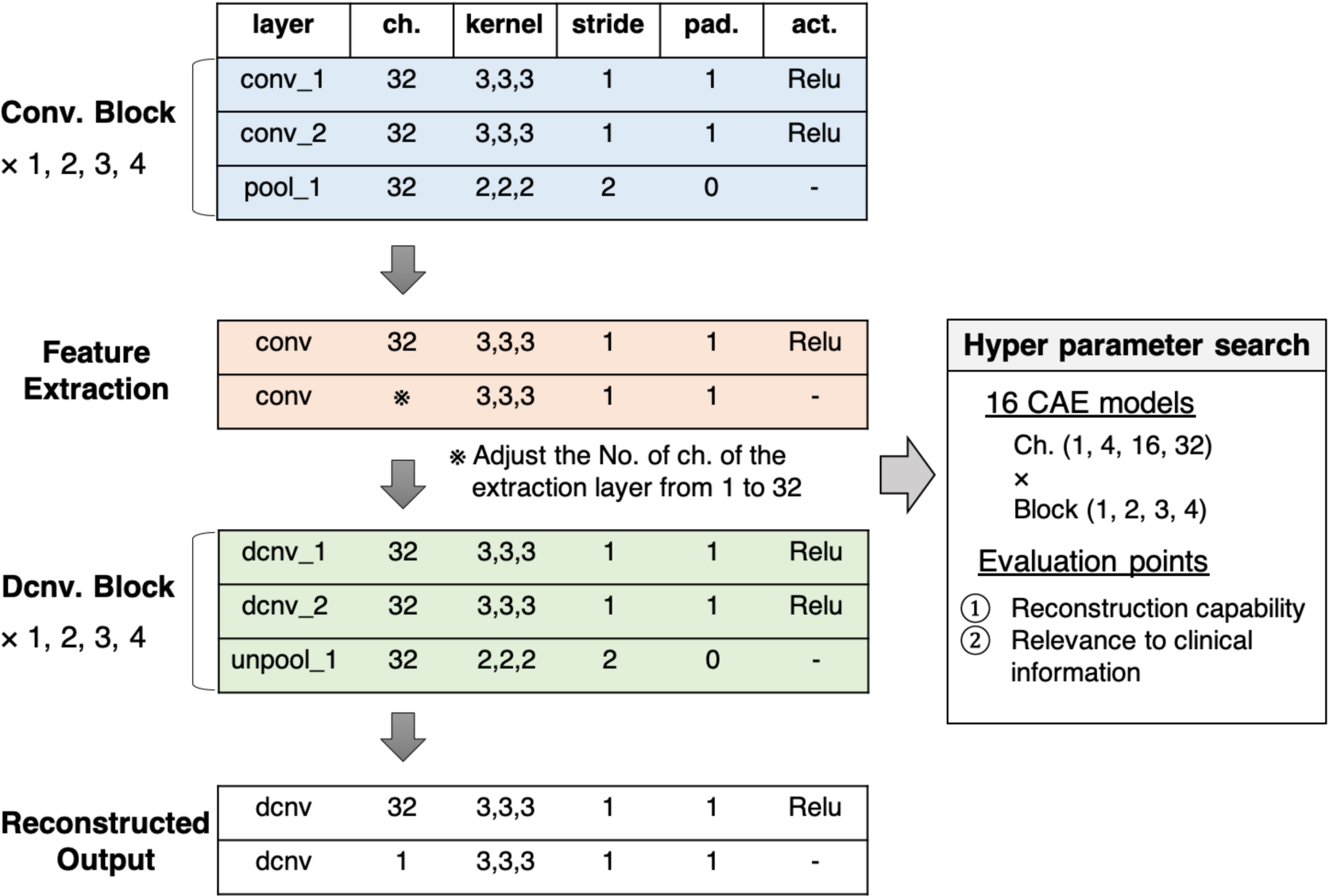
Our proposed CAE architecture. One convolution/deconvolution block was defined as repeating two convolution/deconvolution layers and one pooling/unpooling layer. The number of blocks was set from 1 to 4. The number of channels in the extraction layer was set from 1 to 32. Sixteen patterns of models with different numbers of blocks and channels were developed. In order to explore the effective number of channels and blocks, the reconstruction capability and relevance to clinical information were evaluated. act., activation function; CAE, convolutional autoencoder; ch., channel; Conv., convolution; Dcnv, deconvolution; pad., padding; pool, pooling; Relu, Rectified Linear Unit; unpool, unpooling.

Other hyper parameters were fixed and common among models. The encoder was composed of convolution layers (a kernel size of 3×3×3 and a stride of 1) with rectified linear unit (ReLU) activations and average pooling layers (a kernel size of 2×2×2 and a stride of 2). The decoder was composed of convolution layers (a kernel size of 3×3×3 and a stride of 1) with ReLU activations and unpooling layers (a kernel size of 2×2×2 and a stride of 2). The loss function consisted of the mean absolute error (MAE) between the input and the reconstruction. As an optimizer, we used a gradient-based method with adaptative learning rates called Adam^42^ (alpha = 0.0001, beta1 = 0.9, beta2 = 0.999) using mini-batches with a size of eight samples. The training process was performed with a maximum 50,000 training iterations. We conducted the experiments in Python 3.6 (https://www.python.org/) using the Chainer v.5.4.0 library^43^.

We used a reference of training performances of 3D-CAEs, referred to as the “average brain”, with which the model was assumed to output the average intensities of the training dataset regardless of the inputs. The average brain is one of the most trivial solutions where the network outputs an image without learning any information about individual differences of the inputs. The signal intensities of voxel *i* of the average brain was determined as follows:

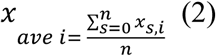

where *s* is a sample from training dataset and *n* is the number of samples.

### Regression analysis with demographic and clinical information

Whether the extracted features retained information relevant to demographic and clinical information was evaluated using linear regression analysis, in which demographic and clinical information were predicted as an objective variable and extracted features were used as explanatory variables (see the lower part of Figure 1). Demographic and clinical information included age, scores of positive and negative symptoms (PANSS), dose of antipsychotic medications (chlorpromazine equivalent [CPZE]), Wechsler Adult Intelligence Scale (WAIS), duration of illness, and age at onset. For the regression analysis, in order to reduce the effects of correlated variables we adopted ridge regression, one of regularized linear regression methods. In the regression analysis, we executed a five-fold cross-validation process whereby the COBRE dataset was randomly divided into five group of samples (folds), and then samples from four folds were used for training the regression model, and the other fold was used for the test of the regression model. The five-fold cross-validation was repeated ten times. Performance of the regression model was evaluated using the root mean square error (RMSE).

Differences in the performances of regression models were evaluated using the two-way (number of channels × number of blocks) analysis of variance (ANOVA). Subsequently, Tukey’s multiple comparison test was performed for each group as a post-hoc analysis. The level of significance was set to 0.05.

The 3D-CAE models were also compared with the ROI method. In the ROI method, using the automated anatomical labeling (AAL) template^11^, the GM was divided into 116 ROIs. The average intensities of each ROI were used as the ROI-based features for regression analysis. Student’s t-test was performed to compare the proposed 3D-CAE model with the ROI method. The level of significance was set to 0.05.

## Results

### Reconstruction capability performance

Figure 3a shows a representative example of learning curves for the 3D-CAE with 16 channels and 3 blocks. Progressive decreases were shown not only with “train loss” (red line), but also “validation loss” (orange line) and “test loss” (green line); this indicated that the 3D-CAE successfully learned to reproduce the high-resolution MRI input data without overfitting. The level of MAEs were remarkably below the level of the “average brain” (dashed line), at which the model is assumed to output the average intensities of the training dataset regardless of the inputs (see Methods for details), suggesting that the 3D-CAE successfully reproduced characteristic features of the individual brains. In addition, the curve for “COBRE loss” (blue line), the reconstruction loss for the images from the COBRE dataset with the model trained by the Kyoto dataset, showed a similar trend. This indicated that the 3D-CAE can be applied to MRI data from another site with different scanning parameters. Similar trends of learning curves were observed for the other fifteen 3D-CAEs with different hyper parameter settings, indicating that all models (sixteen 3D-CAEs with varying hyper parameters) successfully converged to reproduce high resolution MRI input data without overfitting.

**Figure 3.**
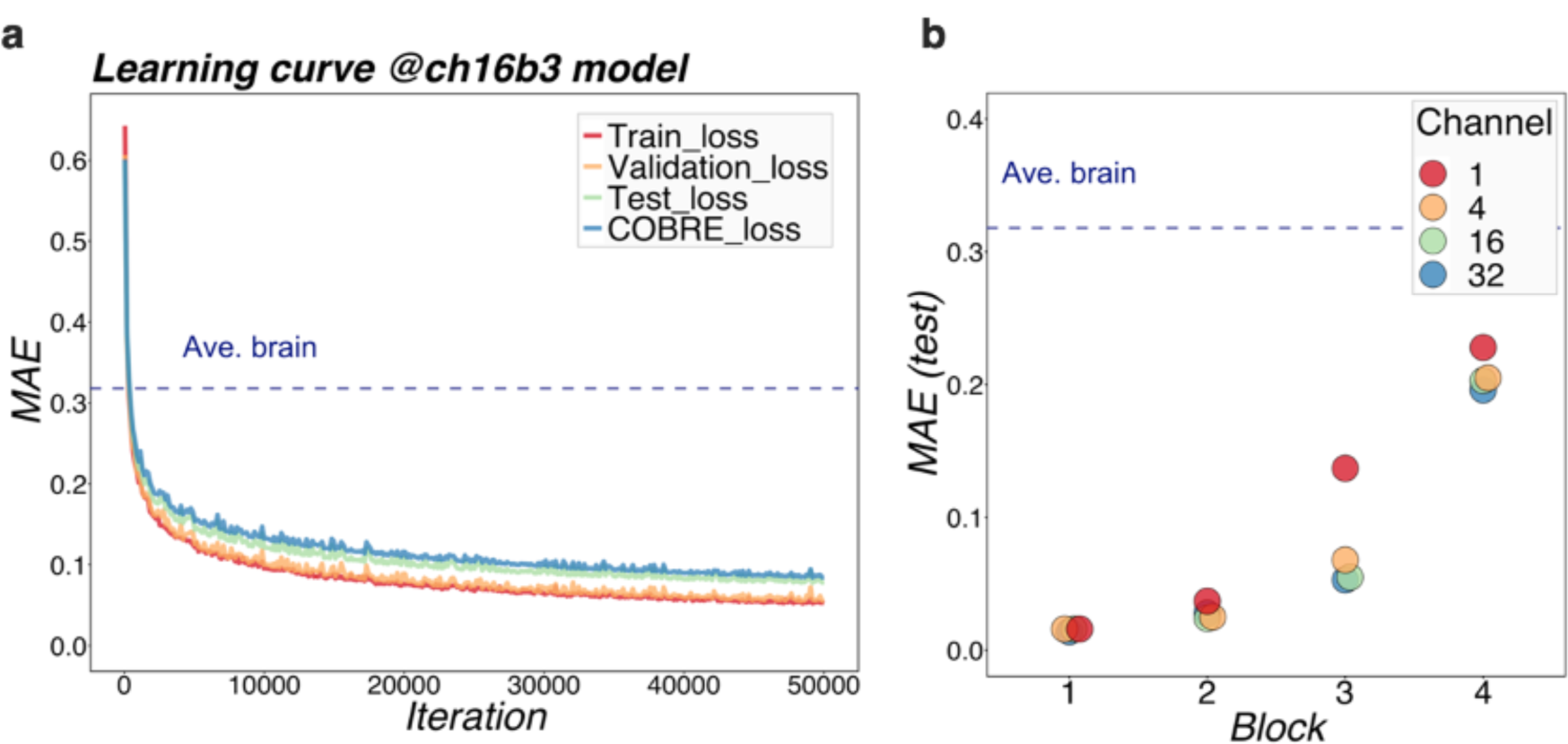
Learning performance of models. (a) shows the learning loss curve for a 16-channel and 3-block model. The red line shows the training loss, indicating that the learning has progressed and the loss has fallen sufficiently. The validation loss and test loss were also decreased, so the model was not overfitting. The blue line indicates the loss at the other site (COBRE), and the loss degraded as well. It can be seen that the MAE of our proposed models was well below the level of Ave.brain at which the model was assumed to output the average brain. This suggested that our 3D-CAE models have successfully reconstructed the brain images with individual characteristics. Similar learning curves were found for other models. In (b), the reconstruction performance of each of the 16 models were compared. The relationships between MAE, number of channels, and number of blocks are shown. The horizontal axis indicates the number of blocks, which is color-coded by the number of channels. As the number of blocks increased, the MAE tended to be larger, and as the number of channels increased, the MAE tended to be slightly smaller. 3D-CAE, three-dimensional convolutional autoencoder; COBRE, Center for Biomedical Research Excellence; MAE, mean absolute error.

Figure 3b summarized the reconstruction performances (MAEs for the COBRE dataset) of the sixteen 3D-CAE models with respect to the number of channels and number of blocks. Regarding the number of blocks, it can be seen that the larger the number of blocks, the larger the reconstruction error. This result is intuitively understandable, in that models with smaller blocks are easier to reconstruct because extracted latent features do not abstract the original image as much (Figure 4). Regarding the number of channels, although the differences were small, there was a tendency for the larger the number of channels to be associated with smaller reconstruction errors (see Table 2 for more details). This result is consistent with the fact that the models with more channels have more expressive capability.

**Table 2.**
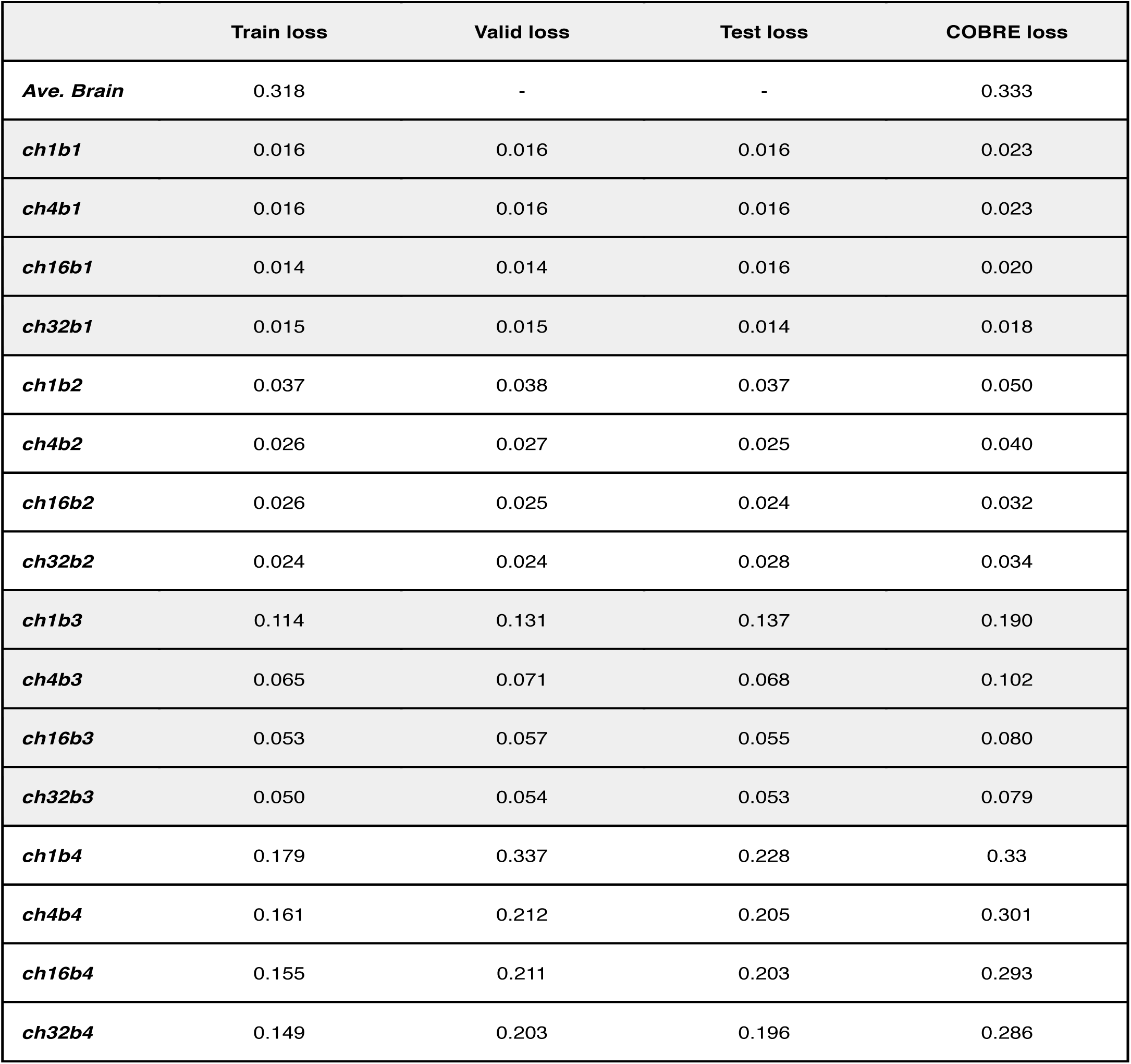
Reconstruction error for each model. Where the ‘Ave. Brain’ of train loss was the average brain in the Kyoto dataset and the ‘Ave. Brain’ of COBRE loss was the average brain in the COBRE dataset.

|  | Train loss | Valid loss | Test loss | COBRE loss |
| --- | --- | --- | --- | --- |
| *Ave. Brain* | 0.318 | - | - | 0.333 |
| *ch1b1* | 0.016 | 0.016 | 0.016 | 0.023 |
| *ch4b1* | 0.016 | 0.016 | 0.016 | 0.023 |
| *ch16b1* | 0.014 | 0.014 | 0.016 | 0.020 |
| *ch32b1* | 0.015 | 0.015 | 0.014 | 0.018 |
| *ch1b2* | 0.037 | 0.038 | 0.037 | 0.050 |
| *ch4b2* | 0.026 | 0.027 | 0.025 | 0.040 |
| *ch16b2* | 0.026 | 0.025 | 0.024 | 0.032 |
| *ch32b2* | 0.024 | 0.024 | 0.028 | 0.034 |
| *ch1b3* | 0.114 | 0.131 | 0.137 | 0.190 |
| *ch4b3* | 0.065 | 0.071 | 0.068 | 0.102 |
| *ch16b3* | 0.053 | 0.057 | 0.055 | 0.080 |
| *ch32b3* | 0.050 | 0.054 | 0.053 | 0.079 |
| *ch1b4* | 0.179 | 0.337 | 0.228 | 0.33 |
| *ch4b4* | 0.161 | 0.212 | 0.205 | 0.301 |
| *ch16b4* | 0.155 | 0.211 | 0.203 | 0.293 |
| *ch32b4* | 0.149 | 0.203 | 0.196 | 0.286 |

**Figure 4:**
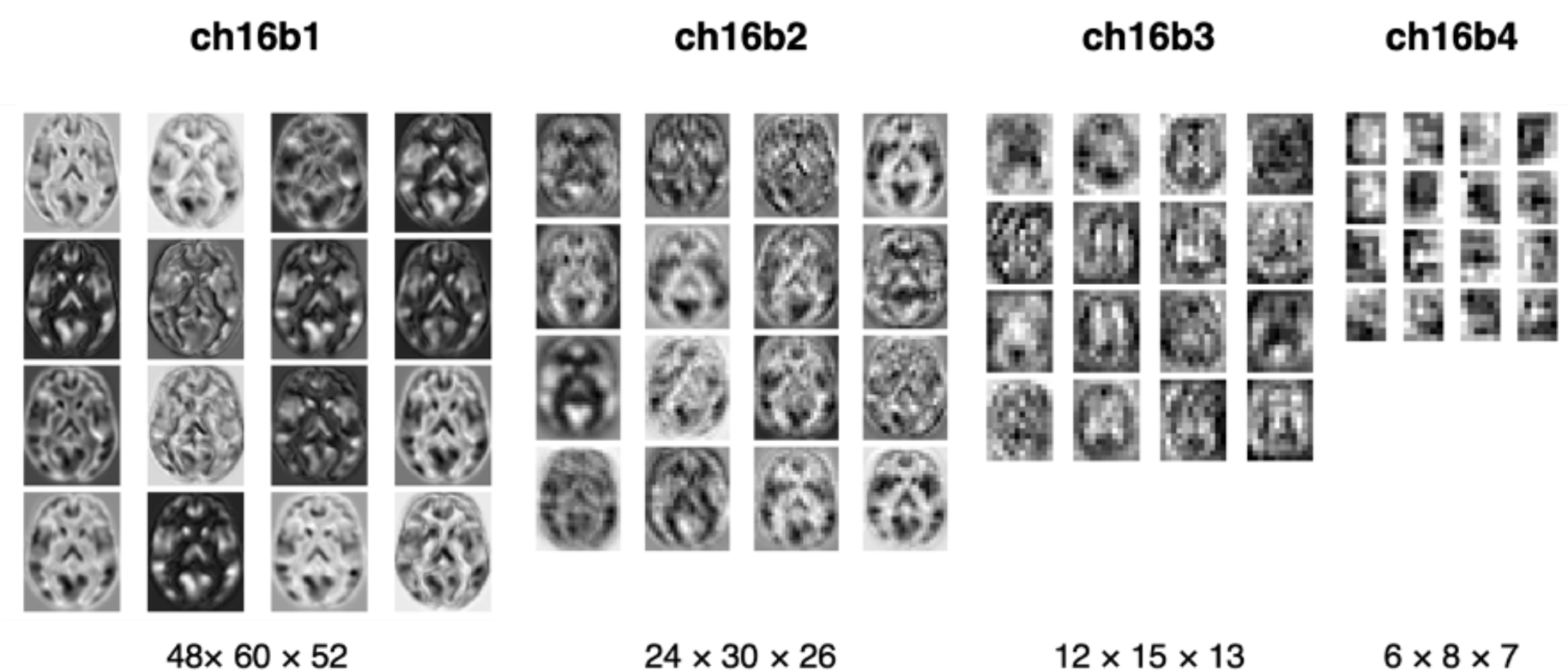
Visualization of extracted features. The extracted features were mapped for four models with 16 channels. From left to right: the model with one, two, three, and four blocks. The middle slices of the horizontal slice from 3D features are shown. In the one-block model, the morphology of the brain can be seen, but with four blocks, the images are more abstract.

### Relevance to clinical information

The efficacy of the proposed method was evaluated using linear regressions for predicting demographic and clinical information related to a psychiatric disorder, i.e., schizophrenia. Demographic and clinical information including age, dose of antipsychotic medication (CPZE), and scores of positive and negative symptoms (PANSS) were used as an objective variable, and all extracted features of 3D-CAE were used as explanatory variables. Features using the ROI-based method were also used for comparison with the conventional method. A linear regression analysis was used as a simplest method to confirm if extracted features from 3D-CAEs with different hyper parameters (numbers of block and channels) preserved useful information. Each of the 16 CAE models were analyzed 10 times, and the difference in predictive performance of the models was examined statistically.

Figure 5 illustrates a representative example of the regression analysis results. Differences in the performance of regression models (RMSE) with respect to the number of channels with 3 blocks (Fig. 5a, b, c, d) and respect to the number of blocks with 16 channels (Fig. 5e, f, g, h) were demonstrated as representative examples. The results of the comparison with the ROI method are shown in Table 3. The detailed results are described in Supplementary Table S1, S2, and S3, respectively.

**Table 3.**
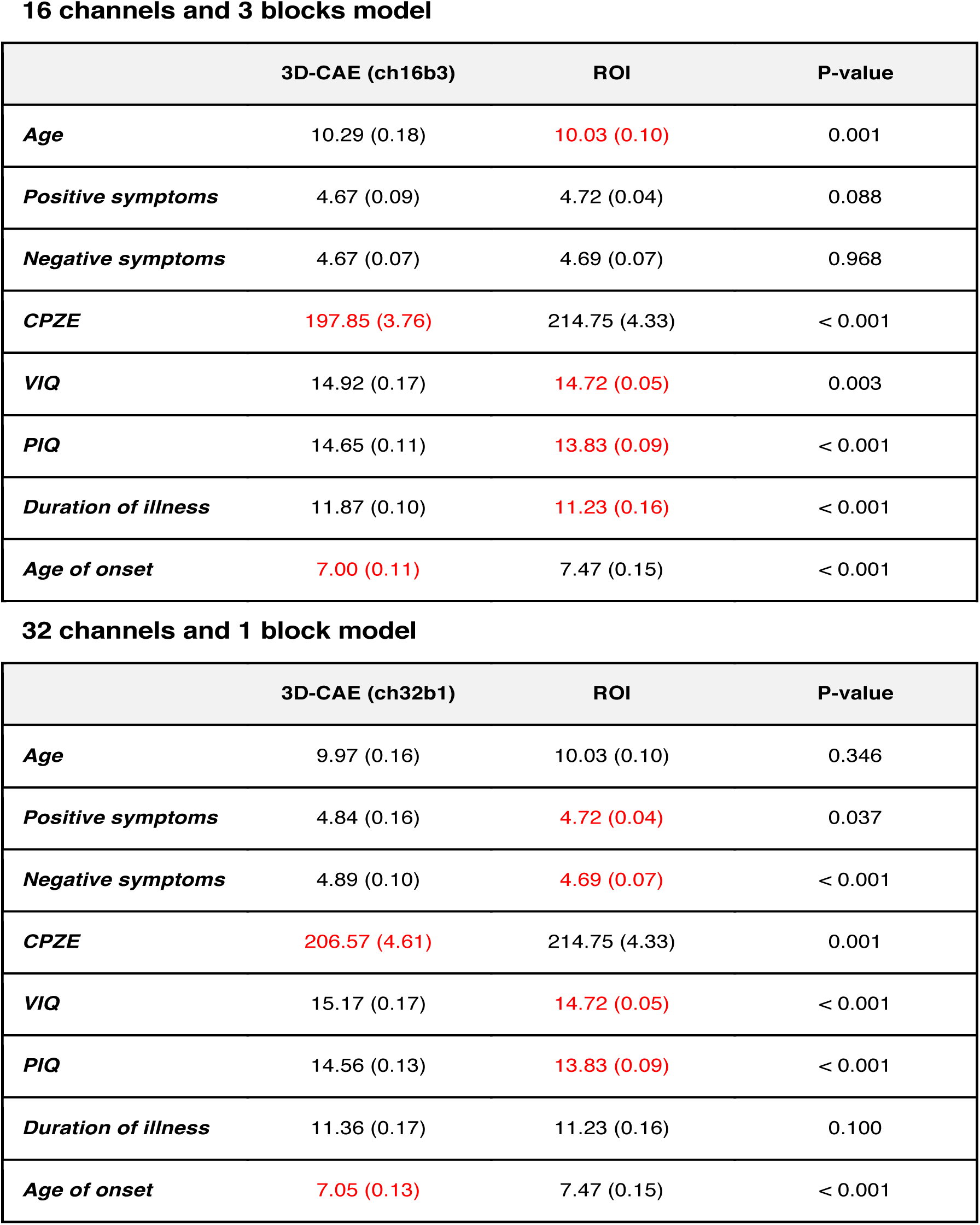
The results of t-test. The differences between 3D-CAE and ROI are shown as mean (standard deviation) and p-value. The significant better performances were marked in red. The 3D-CAE method was superior to the ROI method in predicting CPZE and Age of onset. It seemed that the model with 3 blocks was also comparable or better than the ROI method in Positive symptoms and Negative symptoms.

16 channels and 3 blocks model
|  | 3D-CAE (ch16b3) | ROI | P-value |
| --- | --- | --- | --- |
| **Age** | 10.29 (0.18) | 10.03 (0.10) | 0.001 |
| **Positive symptoms** | 4.67 (0.09) | 4.72 (0.04) | 0.088 |
| **Negative symptoms** | 4.67 (0.07) | 4.69 (0.07) | 0.968 |
| **CPZE** | 197.85 (3.76) | 214.75 (4.33) | < 0.001 |
| **VIQ** | 14.92 (0.17) | 14.72 (0.05) | 0.003 |
| **PIQ** | 14.65 (0.11) | 13.83 (0.09) | < 0.001 |
| **Duration of illness** | 11.87 (0.10) | 11.23 (0.16) | < 0.001 |
| **Age of onset** | 7.00 (0.11) | 7.47 (0.15) | < 0.001 |

**Table 3.** The results of t-test.
|  | 3D-CAE (ch32b1) | ROI | P-value |
| --- | --- | --- | --- |
| **Age** | 9.97 (0.16) | 10.03 (0.10) | 0.346 |
| **Positive symptoms** | 4.84 (0.16) | 4.72 (0.04) | 0.037 |
| **Negative symptoms** | 4.89 (0.10) | 4.69 (0.07) | < 0.001 |
| **CPZE** | 206.57 (4.61) | 214.75 (4.33) | 0.001 |
| **VIQ** | 15.17 (0.17) | 14.72 (0.05) | < 0.001 |
| **PIQ** | 14.56 (0.13) | 13.83 (0.09) | < 0.001 |
| **Duration of illness** | 11.36 (0.17) | 11.23 (0.16) | 0.100 |
| **Age of onset** | 7.05 (0.13) | 7.47 (0.15) | < 0.001 |

**Figure 5.**
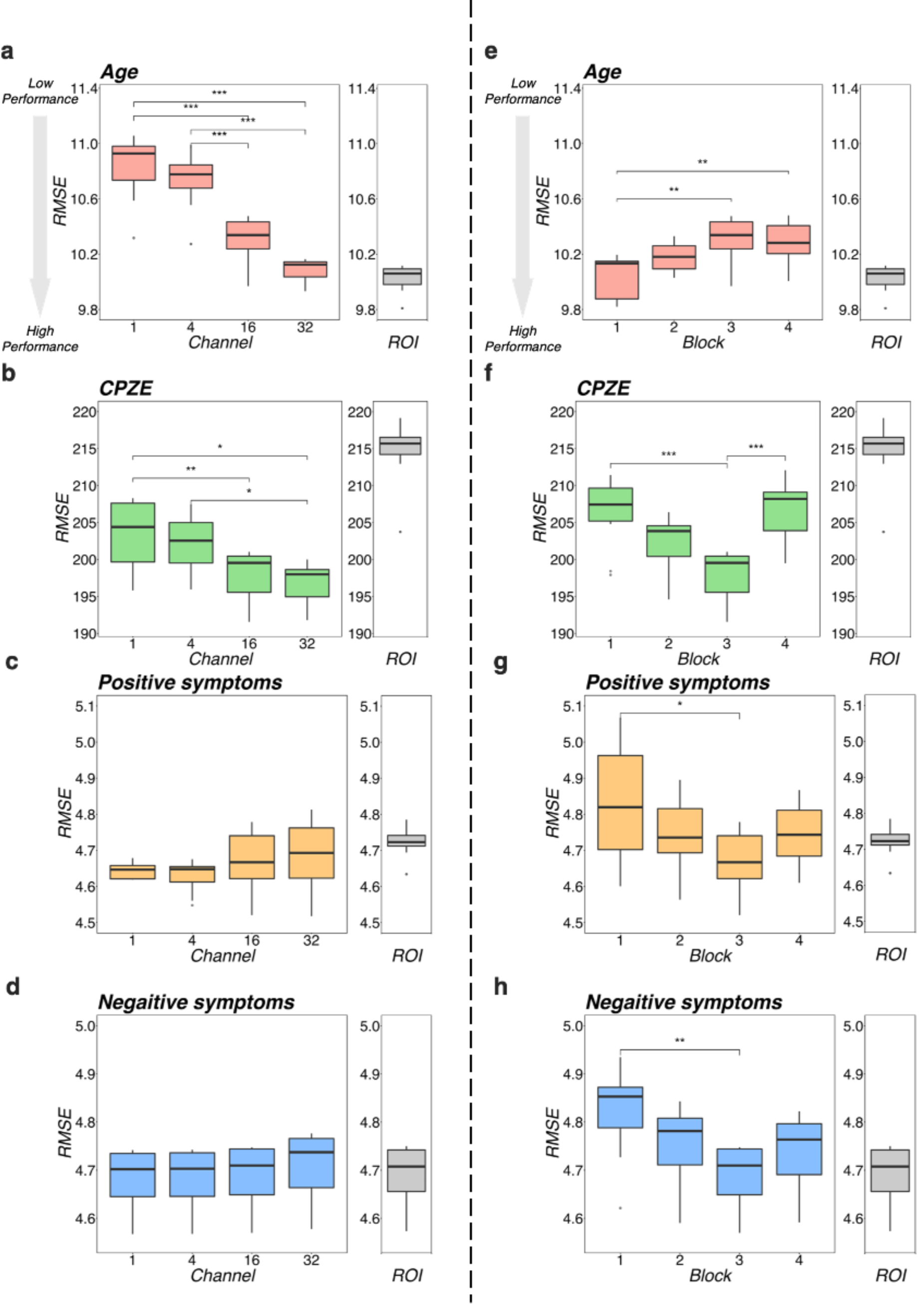
Regression performance plot. The left side (a, b, c, d) shows the model differences by number of channels for the four models with 3 blocks as an example. The right side (e, f, g, h) shows the model differences by number of blocks for the four models, with 16 channels as representative examples. Regarding age, as shown in (a) and (e), the RMSEs were smaller with increasing number of channels and decreasing number of blocks. Regarding CPZE, as shown in (b) and (f), the RMSEs were smaller with increasing number of channels. On the other hand, the RMSEs may be smaller in block 3. Regarding positive symptoms and negative symptoms, as shown in (c) and (d), there was no apparent trend in the number of channels. As shown in (g) and (h), the RMSE may be smaller in block 3. The results of each regression with the ROI method is also included for reference. It suggests that a model with 3 blocks may be appropriate for extracting schizophrenia-related information. *** p <0.001, ** p <0.01, * p <0.05 (two-way analysis of variance followed by Tukey’s multiple comparison test). CPZE, chlorpromazine equivalent; RMSE, root mean square error.

Regarding the prediction of age, there were tendencies for the RMSEs to be smaller with increases in the number of channels (Fig. 5a) and with decreasing number of blocks (Fig. 5b). Indeed, statistical analysis revealed that there were significant differences between the models (channel: p<0.001; block: p<0.001). However, even the model with 32 channels and 1 block, which is considered one of the most predictive models, is equivalent to the ROI method (p = 0.346; Table 3), suggesting that for the prediction of age, 3D-CAE-based features were comparable to a conventional method.

Regarding the prediction for CPZE, there was a tendency for the RMSEs to be smaller with increases in the number of channels (Fig. 5c); on the other hand, the RMSEs were smallest with the condition of 3 blocks (Fig. 5d). Statistical analysis revealed that there were significant differences between the models (channel: p<0.001; block: p<0.001). Post-hoc analysis revealed that there were significant differences between 1 block and 3 blocks, and 3 blocks and 4 blocks. Moreover, the lowest level of RMSE of 3D-CAE was significantly lower than the RMSE from ROI-based features (p<0.001; Table 3), indicating that for the prediction of CPZE, 3D-CAE based features outperformed a conventional method.

Regarding the prediction of positive symptoms, there was no clear tendency with respect to the number of channels (Fig. 5e). On the other hand, with respect to the number of blocks, the RMSEs seemed to be smallest with the condition of 3 blocks (Fig. 5f). Statistical analysis indicated that there were significant differences between the models (channel: p<0.001; block: p<0.001). Post-hoc analysis revealed that there were significant differences between 1 block and 3 blocks. Similar trends could be observed in the prediction of negative symptoms (Fig. 5g, h), where there were significant differences between the models (channel: p<0.001; block: p<0.001). In comparison to the conventional method, although there was no significant difference in the prediction of positive symptoms between the 3D-CAE model with 3 blocks and the ROI method (p=0.088; Table 3), the mean RMSE (SD) was 4.67 (0.09) and 4.72 (0.04), respectively, suggesting that the 3D-CAE might be comparable or better than the ROI method. Regarding the prediction of negative symptoms, there was no significant difference between 3D-CAE and the conventional method (p= 0.968; Table 3).

## Discussion

We have shown that (1) the proposed 3D-CAEs successfully reconstructed high-resolution 3D MRI data with sufficiently low errors, and (2) the diagnostic label-free features extracted using 3D-CAE retained the relevance of various clinical information. In addition, we explored the appropriate hyper parameter range of 3D-CAE and our results suggest that a model with 3 blocks might be relevant to extract features for predicting the medication dose and symptom severity in patients with schizophrenia.

The reconstruction errors of 3D-CAE were lower than the average brain level, indicating that the proposed 3D-CAEs successfully reconstructed high-resolution 3D brain MRI data with individual characteristics. In addition, the 3D-CAE trained with the Kyoto dataset was applicable to the COBRE dataset with different scanners and scanning parameters. Although the current study was tested using only two datasets, the results suggested that the proposed method may have applicability to data from multiple sites and scanners, itself a challenging issue in neuroimaging studies^44-48^.

Regression analyses demonstrated that CAE-based features were efficient to predict medication dose and symptom severity in patients with schizophrenia, even though CAE-based features were extracted without using a diagnostic label of schizophrenia. Moreover, it was comparable or better than the ROI-based features. This suggests that the proposed model may be useful as a method of label-free feature extraction for neuroimaging studies of other psychiatric disorders with heterogeneity problems that are similar to those seen in schizophrenia.

Regarding the number of channels, 16- to 32-channel models demonstrated better performance. This is easy to understand because the more channels the model has, the more expressive it is^1, 39, 49^. However, since increasing number of channels inevitably results in increasing computational power needs, estimation of the appropriate number of channels is still important. Our results suggest that the number of channels may be sufficient at 16 or 32 for reconstructing structural brain MRI scans. Regarding the number of blocks, our results indicated that information from a local receptive field (small number of block) was sufficient for predicting age. On the other hand, prediction of schizophrenia-related clinical data required information from more global receptive fields (larger block numbers, such as 3-block). As the number of blocks increases, the effective receptive fields expand and global features of the brain can be extracted^19, 41, 42, 50^. In our model, the 3 blocks model contained eight convolutional layers, and effective receptive fields of the feature unit were about 68 × 68 × 68 voxels, corresponding to about 30 percent of the brain. This fact is consistent with the previous neuroimaging studies showing that positive symptoms are associated with the volume of multiple brain regions, including the middle temporal gyrus, middle frontal gyrus, and amygdala^51-55^. Similarly, the dose of antipsychotic medication administered has been reported to be associated with the volume of multiple brain regions, including the superior frontal gyrus, medial temporal gyrus, and amygdala^51-54, 56^. The superiority of CAE-based features may be related to the detection of local signal interactions inherent in the convolutional methods; this is in contrast to the ROI-based method, in which information is averaged for each ROI and the local signal interactions are discarded.

There are some limitations to our study. First, the number of dimensions of the features extracted by our proposed model was larger than those of the ROI-based features (116 dimensions). Even the model with the smallest number of dimensions, 1 channel and 4 blocks, had 336 dimensions. In this study, we used a relatively simple network and did not explore complex architecture. However, it may be possible to extract lower-dimensional features without losing the quality of information by elaboration of network architectures. Second, the datasets used in this study only included patients diagnosed with schizophrenia as well as healthy subjects. Considering the heterogeneity of psychiatric disorders, it will be necessary to examine the applicability of diagnostic label-free feature extraction using 3D-CAE to other psychiatric disorders in the future. Third, this study was not able to show the *best* architecture, due to the limited number of data samples for statistical power and computational costs for exploring a wide range of hyper parameters. Fourth, the biological implications of the current results remained unclear. For example, in predicting age, it is not yet clear why the relevance decreases as the number of blocks increases. In addition, the visualization of highly active neurons using Smooth Grad^57^ or Grad-CAM^58^ may be useful for investigating the correspondence between the extracted features and actual brain regions, although more computing power and time are needed.

In this paper, we presented 3D-CAE-based feature extraction for brain structural imaging of psychiatric disorders. We found that 3D-CAE can extract features relevant to clinical information from high-resolution 3D MRI data without diagnostic labels. Our data suggests that 3D-CAE models with effective hyper parameter settings may be able to extract information related to the medication dose and symptom severity in patients with schizophrenia. Moreover, further investigations should focus on the correspondence between the features extracted by the CAE and the accumulated findings from the conventional neuroimaging studies.

## Acknowledgements

This work was partially supported by JSPS KAKENHI Grant Numbers JP17H00740, JP17H04248, JP17H06039, JP18KT0021, JP18K07597, JP19H04998, JP19K17077, JP20H00001, and JP20H00625, Grant-in-Aid for Scientific Research on Innovative Areas from the MEXT Grant Numbers JP20H05064, Strategic International Brain Science Research Promotion Program by the Japan Agency for Medical Research and Development Grant Number 19dm0307008h0002, JST CREST Grant Number JPMJCR16E2 and A grant from SENSHIN Medical Research Foundation. We also would like to thank Editage (www.editage.com) for English language editing.

## Author Contributions

HY, YH, and YY conceived and designed the research. HY and YH conducted the deep learning experiments and analyzed the data. GS, JM, TM, and HT collected MRI data. HY, YH and YY drafted the manuscript. GS, JM, TM, HT, MH, and AH provided critical revisions. All authors contributed to and have approved the final manuscript.

## Additional Information

Competing Interests: The authors declare no competing financial interests.

---

**Supplementary Table S1:**
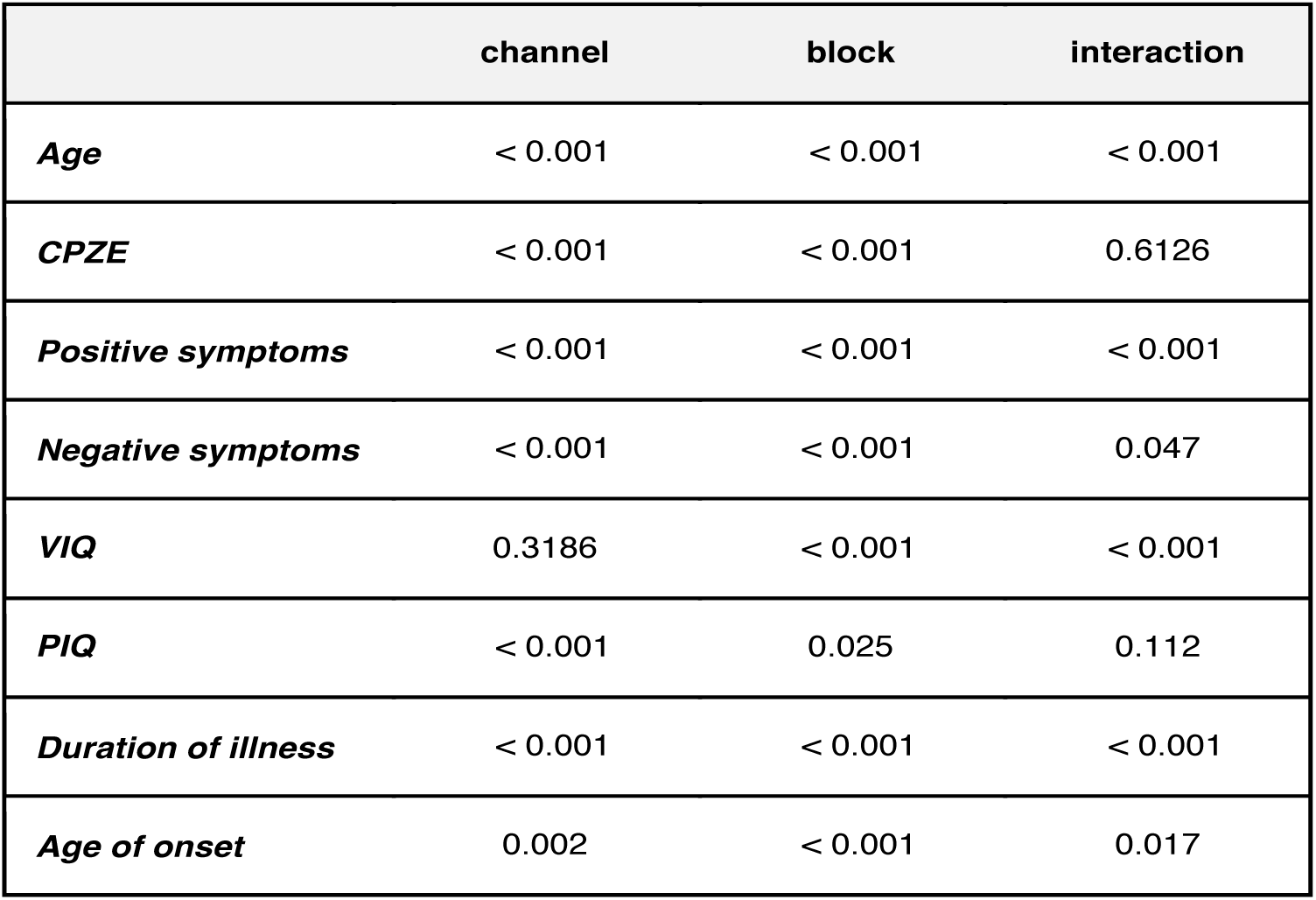
The results of ANOVA. Since there are significant differences in most demographic and clinical information, it can be said that there were differences in the performances depending on the hyper parameters.

|  | channel | block | interaction |
| --- | --- | --- | --- |
| *Age* | < 0.001 | < 0.001 | < 0.001 |
| *CPZE* | < 0.001 | < 0.001 | 0.6126 |
| *Positive symptoms* | < 0.001 | < 0.001 | < 0.001 |
| *Negative symptoms* | < 0.001 | < 0.001 | 0.047 |
| *VIQ* | 0.3186 | < 0.001 | < 0.001 |
| *PIQ* | < 0.001 | 0.025 | 0.112 |
| *Duration of illness* | < 0.001 | < 0.001 | < 0.001 |
| *Age of onset* | 0.002 | < 0.001 | 0.017 |

**Supplementary Table S2:** Regression performance for each model and ROI methods. Listed on Age, CPZE, Positive symptoms, Negative symptoms, VIQ, PIQ, Duration of illness and Age of onset. The average of the regression results was shown. Red ink indicated better performance than that of the ROI method. It seems that that the CAE method was superior to the ROI method in predicting CPZE, Positive symptoms, Negative symptoms, and Age of onset.

|  | Age | CPZE | Positive symptoms | Negative symptoms | VIQ | PIQ | Duration of illness | Age of onset |
| --- | --- | --- | --- | --- | --- | --- | --- | --- |
| **ROI** | 10.03 (0.01) | 214.75 (4.33) | 4.72 (0.04) | 4.69 (0.07) | 14.72 (0.05) | 13.83 (0.09) | 11.23 (0.16) | 7.47 (0.15) |
| **ch1b1** | 10.08 (0.16) | 208.79 (4.54) | 4.67 (0.03) | 4.69 (0.07) | 14.92 (0.08) | 14.59 (0.12) | 11.43 (0.12) | 7.04 (0.12) |
| **ch4b1** | 10.21 (0.22) | 206.94 (4.48) | 4.66 (0.04) | 4.70 (0.07) | 14.91 (0.06) | 14.69 (0.15) | 11.46 (0.13) | 7.10 (0.10) |
| **ch16b1** | 10.04 (0.16) | 206.40 (4.82) | 4.83 (0.16) | 4.82 (0.09) | 14.98 (0.12) | 14.56 (0.11) | 11.43 (0.16) | 7.08 (0.12) |
| **ch32b1** | 9.97 (0.16) | 206.57 (4.61) | 4.84 (0.16) | 4.89 (0.10) | 15.17 (0.17) | 14.56 (0.13) | 11.35 (0.17) | 7.05 (0.13) |
| **ch1b2** | 10.38 (0.17) | 208.14 (4.28) | 4.66 (0.04) | 4.68 (0.07) | 14.87 (0.07) | 14.67 (0.13) | 11.95 (0.13) | 7.02 (0.17) |
| **ch4b2** | 10.42 (0.14) | 203.72 (4.21) | 4.64 (0.05) | 4.69 (0.07) | 14.92 (0.04) | 14.61 (0.10) | 11.71 (0.15) | 7.01 (0.12) |
| **ch16b2** | 10.18 (0.10) | 201.98 (4.12) | 4.75 (0.10) | 4.75 (0.08) | 14.88 (0.10) | 14.55 (0.09) | 11.60 (0.14) | 6.96 (0.10) |
| **ch32b2** | 10.06 (0.10) | 201.01 (4.12) | 4.77 (0.12) | 4.79 (0.09) | 14.86 (0.11) | 14.52 (0.09) | 11.59 (0.14) | 6.89 (0.12) |
| **ch1b3** | 10.83 (0.23) | 203.50 (4.60) | 4.64 (0.02) | 4.68 (0.07) | 14.83 (0.08) | 14.73 (0.12) | 11.94 (0.13) | 7.17 (0.16) |
| **ch4b3** | 10.73 (0.21) | 202.33 (4.07) | 4.63 (0.04) | 4.68 (0.07) | 14.84 (0.11) | 14.70 (0.11) | 12.03 (0.10) | 6.95 (0.10) |
| **ch16b3** | 10.29 (0.18) | 197.85 (3.76) | 4.67 (0.09) | 4.69 (0.07) | 14.91 (0.17) | 14.65 (0.10) | 11.87 (0.10) | 7.00 (0.11) |
| **ch32b3** | 10.09 (0.08) | 196.85 (3.08) | 4.69 (0.10) | 4.71 (0.07) | 14.80 (0.12) | 14.53 (0.09) | 11.78 (0.10) | 6.96 (0.11) |
| **ch1b4** | 10.47 (0.14) | 209.73 (7.02) | 4.65 (0.04) | 4.68 (0.07) | 14.88 (0.07) | 14.75 (0.09) | 11.83 (0.14) | 7.19 (0.21) |
| **ch4b4** | 10.72 (0.17) | 207.91 (3.90) | 4.71 (0.08) | 4.71 (0.07) | 14.86 (0.13) | 14.69 (0.12) | 11.95 (0.13) | 7.31 (0.27) |
| **ch16b4** | 10.28 (0.15) | 207.04 (4.09) | 4.75 (0.09) | 4.74 (0.08) | 14.86 (0.12) | 14.55 (0.12) | 11.92 (0.13) | 7.16 (0.16) |
| **ch32b4** | 10.07 (0.13) | 205.11 (4.55) | 4.75 (0.15) | 4.79 (0.09) | 14.83 (0.15) | 14.53 (0.10) | 11.67 (0.12) | 7.06 (0.08) |

**Supplementary Table S3:**
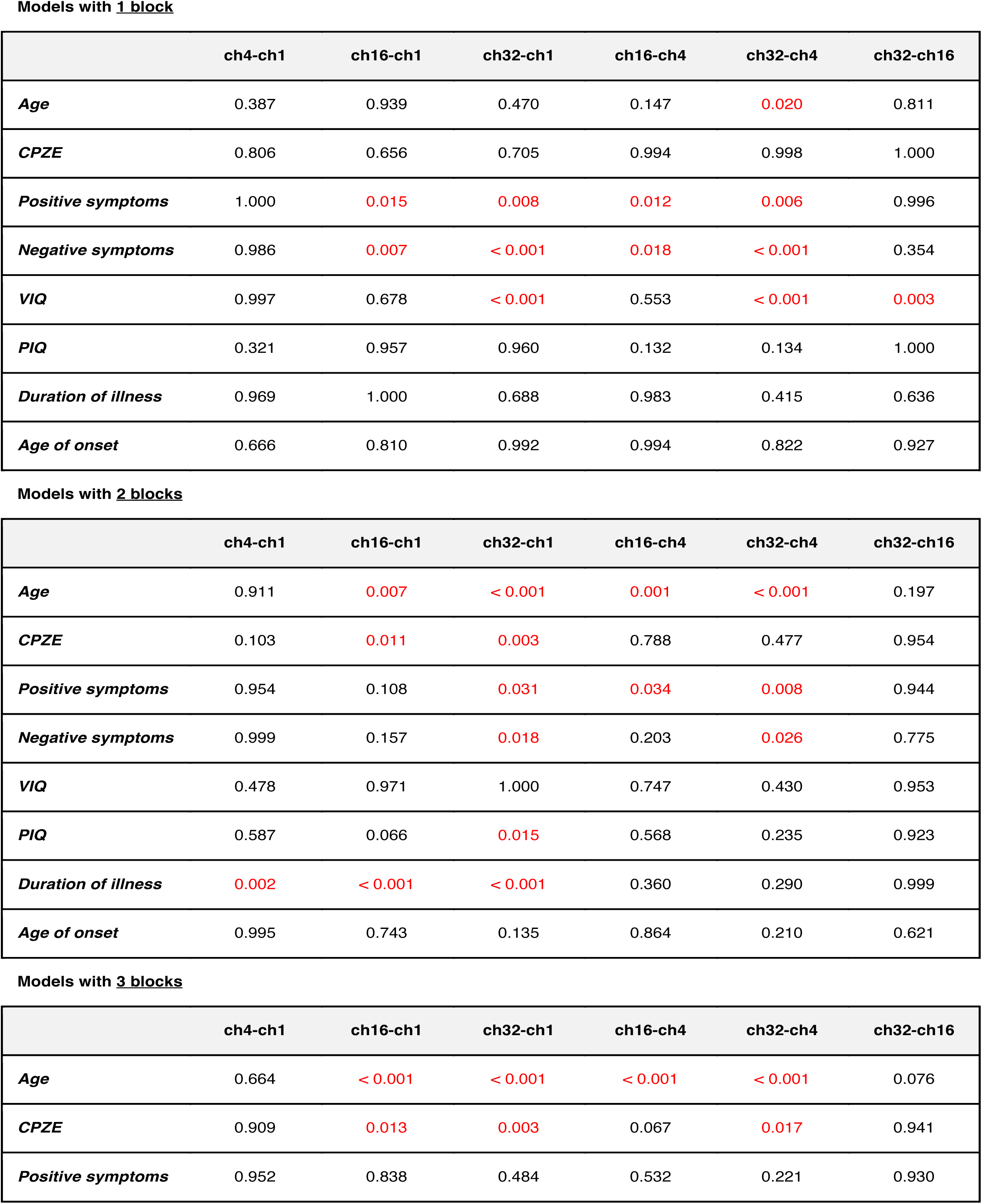
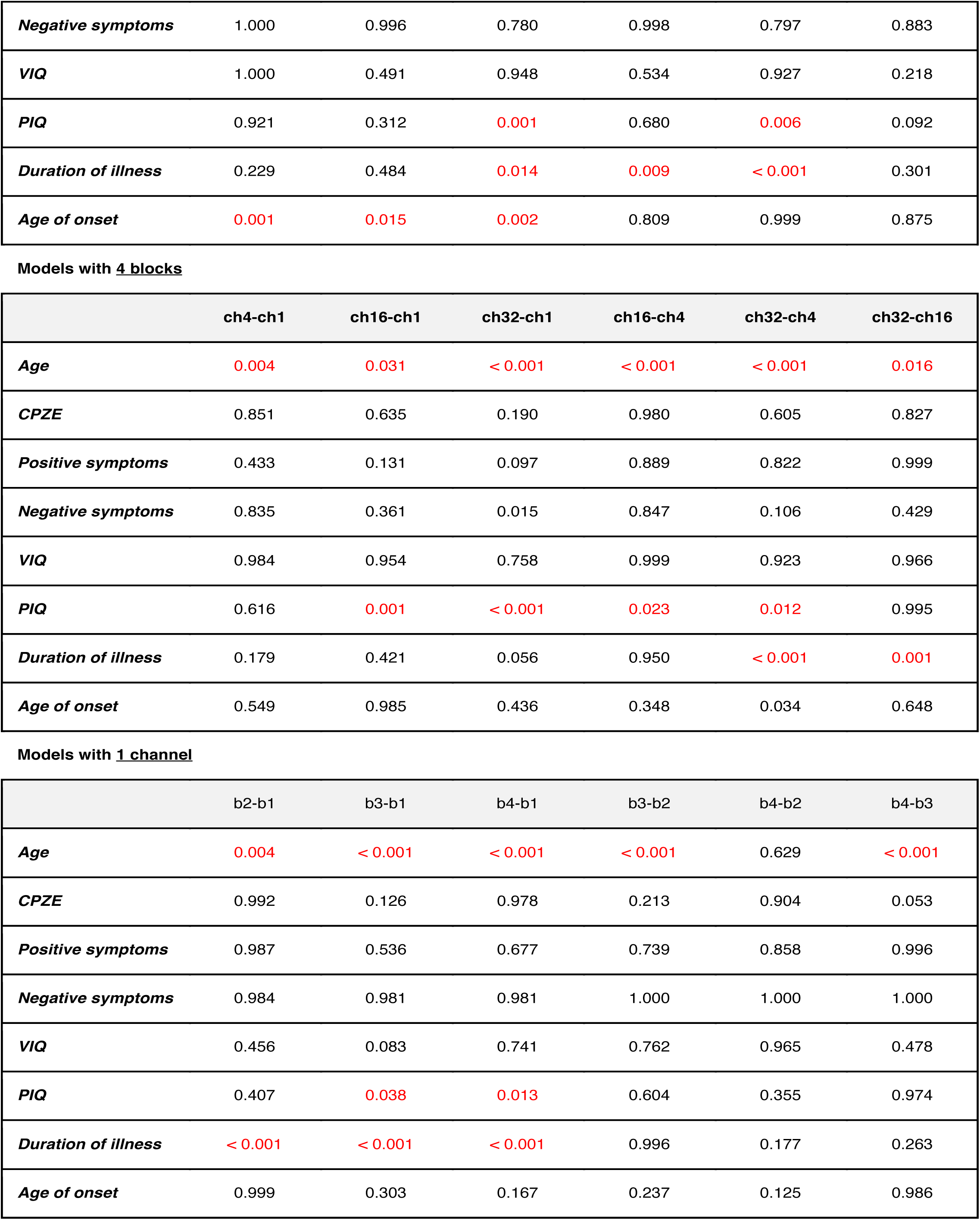
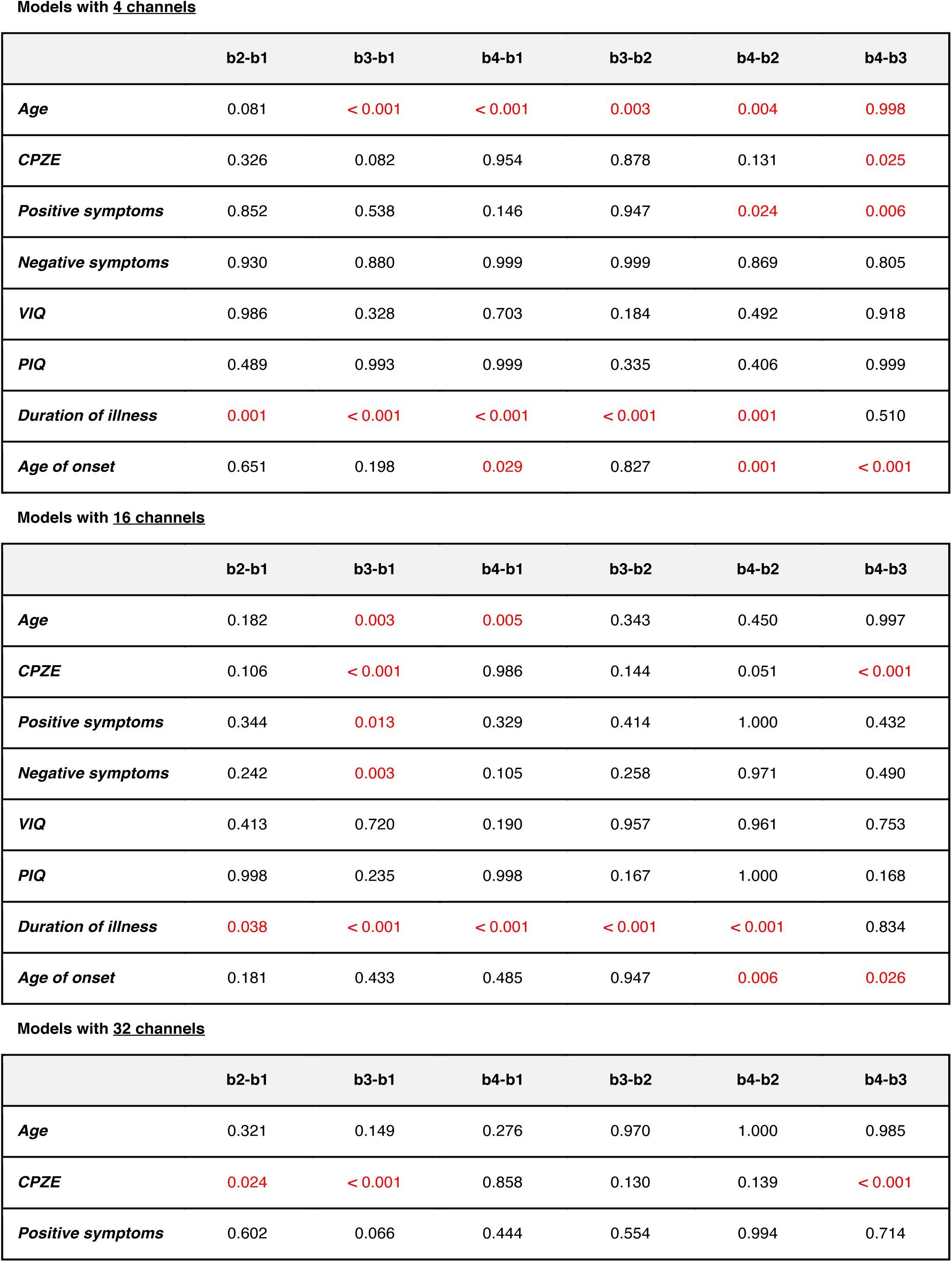
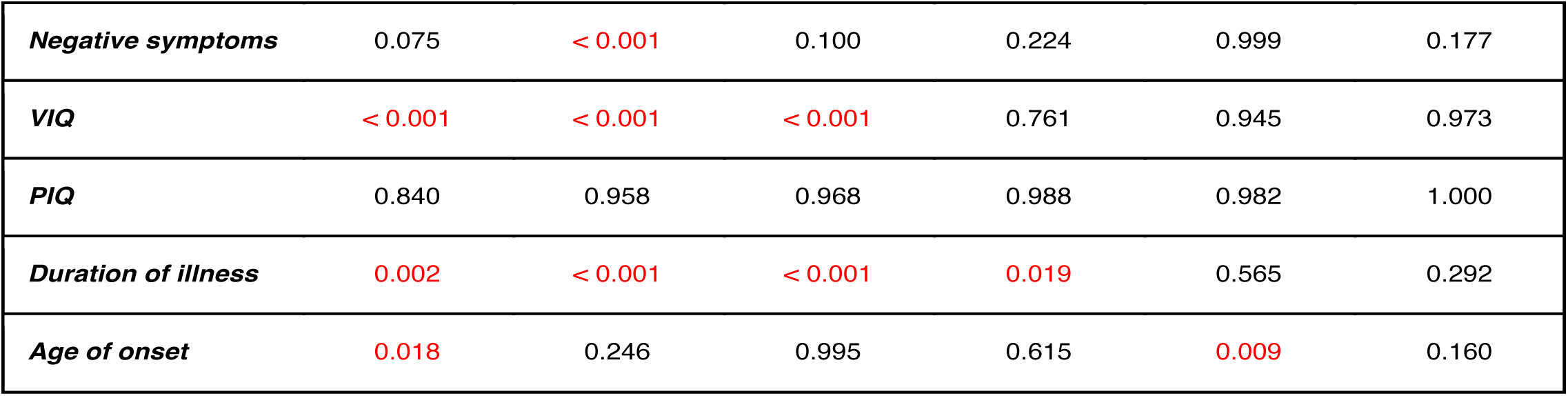
The results of post-hoc analysis. The differences between the different hyper parameter for a model with a particular number of blocks or channels were shown. The numbers listed were p-values and red ink indicated a significant difference.

|  | ch4-ch1 | ch16-ch1 | ch32-ch1 | ch16-ch4 | ch32-ch4 | ch32-ch16 |
| --- | --- | --- | --- | --- | --- | --- |
| *Age* | 0.387 | 0.939 | 0.470 | 0.147 | 0.020 | 0.811 |
| *CPZE* | 0.806 | 0.656 | 0.705 | 0.994 | 0.998 | 1.000 |
| *Positive symptoms* | 1.000 | 0.015 | 0.008 | 0.012 | 0.006 | 0.996 |
| *Negative symptoms* | 0.986 | 0.007 | < 0.001 | 0.018 | < 0.001 | 0.354 |
| *VIQ* | 0.997 | 0.678 | < 0.001 | 0.553 | < 0.001 | 0.003 |
| *PIQ* | 0.321 | 0.957 | 0.960 | 0.132 | 0.134 | 1.000 |
| *Duration of illness* | 0.969 | 1.000 | 0.688 | 0.983 | 0.415 | 0.636 |
| *Age of onset* | 0.666 | 0.810 | 0.992 | 0.994 | 0.822 | 0.927 |

Models with 2 blocks
|  | ch4-ch1 | ch16-ch1 | ch32-ch1 | ch16-ch4 | ch32-ch4 | ch32-ch16 |
| --- | --- | --- | --- | --- | --- | --- |
| *Age* | 0.911 | 0.007 | < 0.001 | 0.001 | < 0.001 | 0.197 |
| *CPZE* | 0.103 | 0.011 | 0.003 | 0.788 | 0.477 | 0.954 |
| *Positive symptoms* | 0.954 | 0.108 | 0.031 | 0.034 | 0.008 | 0.944 |
| *Negative symptoms* | 0.999 | 0.157 | 0.018 | 0.203 | 0.026 | 0.775 |
| *VIQ* | 0.478 | 0.971 | 1.000 | 0.747 | 0.430 | 0.953 |
| *PIQ* | 0.587 | 0.066 | 0.015 | 0.568 | 0.235 | 0.923 |
| *Duration of illness* | 0.002 | < 0.001 | < 0.001 | 0.360 | 0.290 | 0.999 |
| *Age of onset* | 0.995 | 0.743 | 0.135 | 0.864 | 0.210 | 0.621 |

Models with 3 blocks
|  | ch4-ch1 | ch16-ch1 | ch32-ch1 | ch16-ch4 | ch32-ch4 | ch32-ch16 |
| --- | --- | --- | --- | --- | --- | --- |
| *Age* | 0.664 | < 0.001 | < 0.001 | < 0.001 | < 0.001 | 0.076 |
| *CPZE* | 0.909 | 0.013 | 0.003 | 0.067 | 0.017 | 0.941 |
| *Positive symptoms* | 0.952 | 0.838 | 0.484 | 0.532 | 0.221 | 0.930 |
| **Negative symptoms** | 1.000 | 0.996 | 0.780 | 0.998 | 0.797 | 0.883 |
| **VIQ** | 1.000 | 0.491 | 0.948 | 0.534 | 0.927 | 0.218 |
| **PIQ** | 0.921 | 0.312 | 0.001 | 0.680 | 0.006 | 0.092 |
| **Duration of illness** | 0.229 | 0.484 | 0.014 | 0.009 | < 0.001 | 0.301 |
| **Age of onset** | 0.001 | 0.015 | 0.002 | 0.809 | 0.999 | 0.875 |

Models with 4 blocks
|  | ch4-ch1 | ch16-ch1 | ch32-ch1 | ch16-ch4 | ch32-ch4 | ch32-ch16 |
| --- | --- | --- | --- | --- | --- | --- |
| **Age** | 0.004 | 0.031 | < 0.001 | < 0.001 | < 0.001 | 0.016 |
| **CPZE** | 0.851 | 0.635 | 0.190 | 0.980 | 0.605 | 0.827 |
| **Positive symptoms** | 0.433 | 0.131 | 0.097 | 0.889 | 0.822 | 0.999 |
| **Negative symptoms** | 0.835 | 0.361 | 0.015 | 0.847 | 0.106 | 0.429 |
| **VIQ** | 0.984 | 0.954 | 0.758 | 0.999 | 0.923 | 0.966 |
| **PIQ** | 0.616 | 0.001 | < 0.001 | 0.023 | 0.012 | 0.995 |
| **Duration of illness** | 0.179 | 0.421 | 0.056 | 0.950 | < 0.001 | 0.001 |
| **Age of onset** | 0.549 | 0.985 | 0.436 | 0.348 | 0.034 | 0.648 |

Models with 1 channel
|  | b2-b1 | b3-b1 | b4-b1 | b3-b2 | b4-b2 | b4-b3 |
| --- | --- | --- | --- | --- | --- | --- |
| **Age** | 0.004 | < 0.001 | < 0.001 | < 0.001 | 0.629 | < 0.001 |
| **CPZE** | 0.992 | 0.126 | 0.978 | 0.213 | 0.904 | 0.053 |
| **Positive symptoms** | 0.987 | 0.536 | 0.677 | 0.739 | 0.858 | 0.996 |
| **Negative symptoms** | 0.984 | 0.981 | 0.981 | 1.000 | 1.000 | 1.000 |
| **VIQ** | 0.456 | 0.083 | 0.741 | 0.762 | 0.965 | 0.478 |
| **PIQ** | 0.407 | 0.038 | 0.013 | 0.604 | 0.355 | 0.974 |
| **Duration of illness** | < 0.001 | < 0.001 | < 0.001 | 0.996 | 0.177 | 0.263 |
| **Age of onset** | 0.999 | 0.303 | 0.167 | 0.237 | 0.125 | 0.986 |

Models with 4 channels
|  | **b2-b1** | **b3-b1** | **b4-b1** | **b3-b2** | **b4-b2** | **b4-b3** |
| --- | --- | --- | --- | --- | --- | --- |
| *Age* | 0.081 | < 0.001 | < 0.001 | 0.003 | 0.004 | 0.998 |
| *CPZE* | 0.326 | 0.082 | 0.954 | 0.878 | 0.131 | 0.025 |
| *Positive symptoms* | 0.852 | 0.538 | 0.146 | 0.947 | 0.024 | 0.006 |
| *Negative symptoms* | 0.930 | 0.880 | 0.999 | 0.999 | 0.869 | 0.805 |
| *VIQ* | 0.986 | 0.328 | 0.703 | 0.184 | 0.492 | 0.918 |
| *PIQ* | 0.489 | 0.993 | 0.999 | 0.335 | 0.406 | 0.999 |
| *Duration of illness* | 0.001 | < 0.001 | < 0.001 | < 0.001 | 0.001 | 0.510 |
| *Age of onset* | 0.651 | 0.198 | 0.029 | 0.827 | 0.001 | < 0.001 |

Models with 16 channels
|  | **b2-b1** | **b3-b1** | **b4-b1** | **b3-b2** | **b4-b2** | **b4-b3** |
| --- | --- | --- | --- | --- | --- | --- |
| *Age* | 0.182 | 0.003 | 0.005 | 0.343 | 0.450 | 0.997 |
| *CPZE* | 0.106 | < 0.001 | 0.986 | 0.144 | 0.051 | < 0.001 |
| *Positive symptoms* | 0.344 | 0.013 | 0.329 | 0.414 | 1.000 | 0.432 |
| *Negative symptoms* | 0.242 | 0.003 | 0.105 | 0.258 | 0.971 | 0.490 |
| *VIQ* | 0.413 | 0.720 | 0.190 | 0.957 | 0.961 | 0.753 |
| *PIQ* | 0.998 | 0.235 | 0.998 | 0.167 | 1.000 | 0.168 |
| *Duration of illness* | 0.038 | < 0.001 | < 0.001 | < 0.001 | < 0.001 | 0.834 |
| *Age of onset* | 0.181 | 0.433 | 0.485 | 0.947 | 0.006 | 0.026 |

|  | **b2-b1** | **b3-b1** | **b4-b1** | **b3-b2** | **b4-b2** | **b4-b3** |
| --- | --- | --- | --- | --- | --- | --- |
| *Age* | 0.321 | 0.149 | 0.276 | 0.970 | 1.000 | 0.985 |
| *CPZE* | 0.024 | < 0.001 | 0.858 | 0.130 | 0.139 | < 0.001 |
| *Positive symptoms* | 0.602 | 0.066 | 0.444 | 0.554 | 0.994 | 0.714 |
| ***Negative symptoms*** | 0.075 | < 0.001 | 0.100 | 0.224 | 0.999 | 0.177 |
| ***VIQ*** | < 0.001 | < 0.001 | < 0.001 | 0.761 | 0.945 | 0.973 |
| ***PIQ*** | 0.840 | 0.958 | 0.968 | 0.988 | 0.982 | 1.000 |
| ***Duration of illness*** | 0.002 | < 0.001 | < 0.001 | 0.019 | 0.565 | 0.292 |
| ***Age of onset*** | 0.018 | 0.246 | 0.995 | 0.615 | 0.009 | 0.160 |

## Notes

### Competing Interest Statement

The authors have declared no competing interest.

## References

1. Lecun, Y., Bengio, Y. & Hinton, G. Deep learning. Nature 521, 436–444 (2015).

2. Esteva, A. et al. A guide to deep learning in healthcare. Nature Medicine 25, 24–29 (2019).

3. Olesen, J., Gustavsson, A., Svensson, M., Wittchen, H. U. & Jönsson, B. The economic cost of brain disorders in Europe. Eur. J. Neurol. 19, 155–162 (2012).

4. Whiteford, H. A. et al. Global burden of disease attributable to mental and substance use disorders: Findings from the global burden of disease study 2010. Lancet 382, 1575–1586 (2013).

5. Vieira, S., Pinaya, W. H. L. & Mechelli, A. Using deep learning to investigate the neuroimaging correlates of psychiatric and neurological disorders: Methods and applications. Neurosci. Biobehav. Rev. 74, 58–75 (2017).

6. Feczko, E. et al. The heterogeneity problem: Approaches to identify psychiatric subtypes. Trends Cogn. Sci. 23, 584–601 (2019).

7. Linden, D. E. J. The challenges and promise of neuroimaging in psychiatry. Neuron 73, 8–22 (2012).

8. Fornito, A., Zalesky, A., Pantelis, C. & Bullmore, E. T. Schizophrenia, neuroimaging and connectomics. NeuroImage 62, 2296–2314 (2012).

9. Fusar-Poli, P., Howes, O., Bechdolf, A. & Borgwardt, S. Mapping vulnerability to bipolar disorder: A systematic review and meta-analysis of neuroimaging studies. J. Psychiatry Neurosci. 37, 170–184 (2012).

10. Ratnanather, J. T. et al. Morphometry of superior temporal gyrus and planum temporale in schizophrenia and psychotic bipolar disorder. Schizophr. Res. 150, 476–483 (2013).

11. Tzourio-Mazoyer, N. et al. Automated anatomical labeling of activations in SPM using a macroscopic anatomical parcellation of the MNI MRI single-subject brain. NeuroImage 15, 273–289 (2002).

12. Desikan, R. S. et al. An automated labeling system for subdividing the human cerebral cortex on MRI scans into gyral based regions of interest. Neuroimage 31, 968–980 (2006).

13. Nelson, B. G., Bassett, D. S., Camchong, J., Bullmore, E. T. & Lim, K. O. Comparison of large-scale human brain functional and anatomical networks in schizophrenia. NeuroImage Clin. 15, 439–448 (2017).

14. Poldrack, R. A. Region of interest analysis for fMRI. Soc. Cogn. Affect. Neurosci. 2, 67–70 (2007).

15. Heinsfeld, A. S., Franco, A. R., Craddock, R. C., Buchweitz, A. & Meneguzzi, F. Identification of autism spectrum disorder using deep learning and the ABIDE dataset. NeuroImage Clin. 17, 16–23 (2018).

16. Pinaya, W. H. L., Mechelli, A. & Sato, J. R. Using deep autoencoders to identify abnormal brain structural patterns in neuropsychiatric disorders: A large-scale multi-sample study. Hum. Brain Mapp. 40, 944–954 (2019).

17. Aghdam, M. A., Sharifi, A. & Pedram, M. M. Diagnosis of autism spectrum disorders in young children based on resting-state functional magnetic resonance imaging data using convolutional neural networks. J. Digit. Imaging 32, 899–918 (2019).

18. Sarraf, S., DeSouza, D. D., Anderson, J., Tofighi, G. & Initiativ, for the A. D. N. DeepAD: Alzheimer’s disease classification via deep convolutional neural networks using MRI and fMRI. bioRxiv 070441 (2017). doi: 10.1101/070441

19. Wang, S. H. et al. Classification of alzheimer’s disease based on eight-layer convolutional neural network with leaky rectified linear unit and max pooling. J. Med. Syst. 42, 85 (2018).

20. Qureshi, M. N. I., Oh, J. & Lee, B. 3D-CNN based discrimination of schizophrenia using resting-state fMRI. Artif. Intell. Med. 98, 10–17 (2019).

21. Wang, Z., Sun, Y., Shen, Q. & Cao, L. Dilated 3D convolutional neural networks for brain MRI data classification. IEEE Access 7, 134388–134398 (2019).

22. Association, A. P. Diagnostic and statistical manual of mental disorders (DSM-5®). (American Psychiatric Pub, 2013).

23. Organization., W. H. International statistical classification of diseases and related health problems. (WHO, 1992).

24. Owen, M. J. New approaches to psychiatric diagnostic classification. Neuron 84, 564–571 (2014).

25. Lee, S. et al. Genetic relationship between five psychiatric disorders estimated from genome-wide SNPs Cross-Disorder Group of the Psychiatric Genomics Consortium. Nat. Genet. 45, 984–994 (2014).

26. van Os, J. & Kapur, S. Schizophrenia. Lancet 374, 635–645 (2009).

27. Owen, M. J., Sawa, A. & Mortensen, P. B. Schizophrenia. Lancet 388, 86–97 (2016).

28. Plis, S. M. et al. Deep learning for neuroimaging: a validation study. Front. Neurosci. 8, 1–11 (2014).

29. Ladjal, S., Newson, A., & Pham, C. A PCA-like Autoencoder. Preprint at https://arxiv.org/abs/1904.01277 (2019).

30. Martinez-Murcia, F. J., Ortiz, A., Gorriz, J. M., Ramirez, J. & Castillo-Barnes, D. Studying the manifold structure of alzheimer’s disease: A deep learning approach using convolutional autoencoders. IEEE J. Biomed. Heal. Informatics 24, 17–26 (2020).

31. Sugihara, G. et al. Distinct patterns of cerebral cortical thinning in schizophrenia: A neuroimaging data-driven approach. Schizophr. Bull. 43, 900–906 (2017).

32. Association., A. P. Diagnostic and statistical manual of mental disorders: DMS-IV. (American Psychiatric Pub, 1994).

33. First, M. B. SCID-I: Structured clinical interview for DSM-IV axis I disorders. (American Psychiatric Press, 1997).

34. Kay, S. R., Fiszbein, A. & Opler, L. A. The positive and negative syndrome scale (PANSS) for schizophrenia. Schizophr. Bull. 13, 261–276 (1987).

35. K. Friston, J. Ashburner, S. Kiebel, T. Nichols and W. Penny, Statistical parametric mapping (Academic Press, London, 2007)

36. Ashburner, J. A fast diffeomorphic image registration algorithm. NeuroImage 38, 95–113 (2007).

37. Oh, K. et al. Classification of schizophrenia and normal controls using 3D convolutional neural network and outcome visualization. Schizophr. Res. 212, 186–195 (2019).

38. Nishio, M. et al. Convolutional auto-encoder for image denoising of ultra-low-dose CT. Heliyon 3, 393 (2017).

39. Guo, X., Liu, X., Zhu, E. & Yin, J. Deep clustering with convolutional autoencoders. in Lecture Notes in Computer Science (including subseries Lecture Notes in Artificial Intelligence and Lecture Notes in Bioinformatics) 10635 LNCS, 373–382 (2017).

40. Le, H. & Borji, A. What are the receptive, effective receptive, and projective fields of neurons in convolutional neural networks? Preprint at https://arxiv.org/abs/1904.01277 (2017).

41. Luo, W., Li, Y., Urtasun, R. & Zemel, R. Understanding the effective receptive field in deep convolutional neural networks. Adv. Neural Inf. Process. Syst. 4905–4913 (2017).

42. Kingma, D. P. & Ba, J. L. Adam: A method for stochastic optimization. in 3rd International Conference on Learning Representations, ICLR 2015 - Conference Track Proceedings (2015).

43. Tokui, S., Oono, K., Hido, S. & Clayton, J. Chainer: a next-generation open source framework for deep learning. in Proceedings of workshop on machine learning systems (LearningSys) in the twenty-ninth annual conference on neural information processing systems (NIPS) 5, 1–6 (2015).

44. Jovicich, J. et al. MRI-derived measurements of human subcortical, ventricular and intracranial brain volumes: Reliability effects of scan sessions, acquisition sequences, data analyses, scanner upgrade, scanner vendors and field strengths. NeuroImage 46, 177–192 (2009).

45. Schnack, H. G. et al. Mapping reliability in multicenter MRI: Voxel-based morphometry and cortical thickness. Hum. Brain Mapp. 31, 1967–1982 (2010).

46. Fortin, J. P. et al. Harmonization of cortical thickness measurements across scanners and sites. NeuroImage 167, 104–120 (2018).

47. Yamashita, A. et al. Harmonization of resting-state functional MRI data across multiple imaging sites via the separation of site differences into sampling bias and measurement bias. PLOS Biol. 17, e3000042 (2019).

48. Dewey, B. E. et al. DeepHarmony: A deep learning approach to contrast harmonization across scanner changes. Magn. Reson. Imaging 64, 160–170 (2019).

49. Zhu, H. et al. Rethinking the number of channels for the convolutional neural network. Preprint at https://arxiv.org/abs/1909.01861 (2019).

50. Szegedy, C. et al. Going deeper with convolutions. in Proceedings of the IEEE Computer Society Conference on Computer Vision and Pattern Recognition 07-12-June, 1–9 (IEEE Computer Society, 2015).

51. van Erp, T. G. M. et al. Cortical brain abnormalities in 4474 individuals with schizophrenia and 5098 control subjects via the enhancing neuro imaging genetics through meta analysis (ENIGMA) consortium. Biol. Psychiatry 84, 644–654 (2018).

52. Van Erp, T. G. M. et al. Subcortical brain volume abnormalities in 2028 individuals with schizophrenia and 2540 healthy controls via the ENIGMA consortium. Mol. Psychiatry 21, 547–553 (2016).

53. Fan, F. et al. Subcortical structures and cognitive dysfunction in first episode schizophrenia. Psychiatry Res. Neuroimaging 286, 69–75 (2019).

54. García-Martí, G. et al. Schizophrenia with auditory hallucinations: A voxel-based morphometry study. Prog. Neuro-Psychopharmacology Biol. Psychiatry 32, 72–80 (2008).

55. Bullmore, E. Cortical thickness and connectivity in schizophrenia. Am. J. Psychiatry 176, 505–506 (2019).

56. Palaniyappan, L. et al. Cortical folding defects as markers of poor treatment response in first-episode psychosis. JAMA Psychiatry 70, 1031–1040 (2013).

57. Smilkov, D., Thorat, N., Kim, B., Viégas, F. & Wattenberg, M. SmoothGrad: removing noise by adding noise. Preprint at https://arxiv.org/abs/1706.03825 (2017).

58. Selvaraju, R. R. et al. Grad-CAM: Visual explanations from deep networks via gradient-based localization. in 2017 IEEE International Conference on Computer Vision (ICCV) 2017-Octob, 618–626 (IEEE, 2017).

